# Embodied cognition, embodied regulation, and the Data Rate Theorem

**DOI:** 10.1101/001586

**Authors:** Rodrick Wallace

## Abstract

The Data Rate Theorem carries deep implications for theories of embodied cognition, extensions providing a spectrum of necessary conditions dynamic statistical models useful in empirical studies. A large deviations argument, however, implies that the regulation and stabilization of such systems is itself an interpenetrating phenomenon necessarily convoluted with embodied cognition. For humans, the central regulatory role of culture has long been known. Although a ground-state collapse analogous to generalized anxiety appears ubiquitous to such systems, lack of cultural modulation in real-time automatons or distributed cognition man-machine ‘cockpits’ makes them subject to a pathology under which ‘all possible targets are enemies’.

---

> *…[O]ur first move is simply to treat perception-action problems and language problems as the same kind of thing… Linguistic information is a task resource in exactly the same way as perceptual information… Our behavior emerges from a pool of potential task resources that include the body, the environment and… the brain*.
>
> — *– A.D. Wilson and S. Golonka, 2013*

## 1 Introduction

Varela, Thompson and Rosch (1991), in their study *The Embodied Mind: Cognitive Science and Human Experience*, asserted that the world is portrayed and determined by mutual interaction between the physiology of an organism, its sensimotor circuitry, and the environment. The essential point, in their view, being the inherent structural coupling of brain-body-world. Lively debate has followed and continues (e.g., Clark, 1998; M. Wilson, 2002; A. Wilson and S. Golonka, 2013). See SEP (2011) for details and extensive references. Brooks (1986), and many others, have explored and extended analogous ideas, particularly focusing on robotics. Here, we formalize the basic approach via the Data Rate Theorem, and include as well regulation and stabilization mechanisms, in a unitary construct that must interpenetrate in a similar manner.

Cognition can be described in terms of a sophisticated real-time feedback between interior and exterior, necessarily constrained, as Dretske (1994) has noted, by certain asymptotic limit theorems of probability:

> Communication theory can be interpreted as telling one something important about the conditions that are needed for the transmission of information as ordinarily understood, about what it takes for the transmission of semantic information. This has tempted people… to exploit [information theory] in semantic and cognitive studies…

> …Unless there is a statistically reliable channel of communication between [a source and a receiver]… no signal can carry semantic information… [thus] the channel over which the [semantic] signal arrives [must satisfy] the appropriate statistical constraints of information theory.

Recent intersection of that theory with the formalisms of real-time feedback systems – control theory – may provide insight into matters of embodied cognition and the parallel synergistic problem of embodied regulation and control. Here, we extend that work and apply the resulting conceptual model toward formally characterizing the unitary structural coupling of brain-body-world. In the process, we will explore dynamic statistical models that can be fitted to data.

## 2 The Data-Rate Theorem

The recently-formalized data-rate theorem, a generalization of the classic Bode integral theorem for linear control systems (e.g., Yu and Mehta, 2010; Kitano, 2007; Csete and Doyle, 2002), describes the stability of linear feedback control under data rate constraints (e.g., Mitter, 2001; Tatikonda and Mitter, 2004; Sahai, 2004; Sahai and Mitter, 2006; Minero et al., 2009; Nair et al., 2007; You and Xie, 2013). Given a noise-free data link between a discrete linear plant and its controller, unstable modes can be stabilized only if the feedback data rate ℋ is greater than the rate of ‘topological information’ generated by the unstable system. For the simplest incarnation, if the linear matrix equation of the plant is of the form *x*_*t*+1_ = **A***x_t_* + …, where *x_t_* is the n-dimensional state vector at time *t*, then the necessary condition for stabilizability is

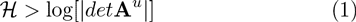

where *det* is the determinant and **A**^*u*^ is the decoupled unstable component of **A**, i.e., the part having eigenvalues ≥ 1.

The essential matter is that there is a critical positive data rate below which there does not exist any quantization and control scheme able to stabilize an unstable (linear) feedback system.

This result, and its variations, are as fundamental as the Shannon Coding and Source Coding Theorems, and the Rate Distortion Theorem (Cover and Thomas, 2006; Ash, 1990; Khinchin, 1957).

We will entertain and extend these considerations, using methods from cognitive theory to explore brain-body-world dynamics that inherently take place under data-rate constraints.

The essential analytic tool will be something much like Pettini’s (2007) ‘topological hypothesis’ – a version of Landau’s spontaneous symmetry breaking insight for physical systems (Landau and Lifshitz, 2007)–which infers that punctuated events often involve a change in the topology of an underlying configuration space, and the observed singularities in the measures of interest can be interpreted as a ‘shadow’ of major topological change happening at a more basic level.

The preferred tool for the study of such topological changes is Morse Theory (Pettini, 2007; Matsumoto, 2002), summarized in the Mathematical Appendix, and we shall construct a relevant Morse Function as a ‘representation’ of the underlying theory.

We begin with recapitulation of an approach to cognition using the asymptotic limit theorems of information theory (Wallace 2000, 2005a, b, 2007, 2012, 2014).

## 3 Cognition as an information source

Atlan and Cohen (1998) argue that the essence of cognition involves comparison of a perceived signal with an internal, learned or inherited picture of the world, and then choice of one response from a much larger repertoire of possible responses. That is, cognitive pattern recognition-and-response proceeds by an algorithmic combination of an incoming external sensory signal with an internal ongoing activity – incorporating the internalized picture of the world – and triggering an appropriate action based on a decision that the pattern of sensory activity requires a response.

Incoming sensory input is thus mixed in an unspecified but systematic manner with internal ongoing activity to create a path of combined signals *x* = (*a*_0_, *a*_1_,…, *a_n_*,…). Each *a_k_* thus represents some functional composition of the internal and the external. An application of this perspective to a standard neural network is given in Wallace (2005a, p.34).

This path is fed into a highly nonlinear, but otherwise similarly unspecified, decision function, *h*, generating an output *h*(*x*) that is an element of one of two disjoint sets *B*_0_ and *B*_1_ of possible system responses. Let

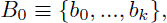

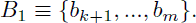

Assume a graded response, supposing that if

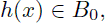

the pattern is not recognized, and if

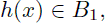

the pattern is recognized, and some action *b_j_*, *k* + 1 ≤ *j* ≤ *m* takes place.

Interest focuses on paths *x* triggering pattern recognition-and-response: given a fixed initial state *a*_0_, examine all possible subsequent paths *x* beginning with *a*_0_ and leading to the event *h*(*x*) ∈ *B*_1_. Thus *h*(*a*_0_,…, *a*_*j*_) ∈ *B*_0_ for all 0 ≤ *j* < *m*, but *h*(*a*_0_, …, *a_m_*) ∈ *B*_1_.

For each positive integer *n*, take *N*(*n*) as the number of high probability paths of length *n* that begin with some particular *a*_0_ and lead to the condition *h*(*x*) ∈ *B*_1_. Call such paths ‘meaningful’, assuming that *N*(*n*) will be considerably less than the number of all possible paths of length *n* leading from *a*_0_ to the condition *h*(*x*) ∈ *B*_1_.

Identification of the ‘alphabet’ of the states *a_j_*, *B_k_* may depend on the proper system coarse graining in the sense of symbolic dynamics (e.g., Beck and Schlogl, 1993).

Combining algorithm, the form of the function *h*, and the details of grammar and syntax, are all unspecified in this model. The assumption permitting inference on necessary conditions constrained by the asymptotic limit theorems of information theory is that the finite limit

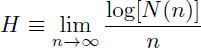

both exists and is independent of the path *x*. Again, *N*(*n*) is the number of high probability paths of length *n*.

Call such a pattern recognition-and-response cognitive process ergodic. Not all cognitive processes are likely to be ergodic, implying that *H*, if it indeed exists at all, is path dependent, although extension to nearly ergodic processes, in a certain sense, seems possible (e.g., Wallace, 2005a, pp. 31-32).

Invoking the Shannon-McMillan Theorem (Cover and Thomas, 2006; Khinchin, 1957), we take it possible to define an adiabatically, piecewise stationary, ergodic information source **X** associated with stochastic variates *X_j_* having joint and conditional probabilities *P*(*a*_0_,…,*a_n_*) and *P*(*a_n_*|*a*_0_,… *a*_*n*−1_) such that appropriate joint and conditional Shannon uncertainties satisfy the classic relations

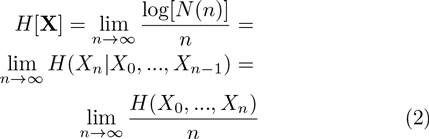

This information source is defined as dual to the underlying ergodic cognitive process, in the sense of Wallace (2005a, 2007).

‘Adiabatic’ means that, when the information source is properly parameterized, within continuous ‘pieces’, changes in parameter values take place slowly enough so that the information source remains as close to stationary and ergodic as needed to make the fundamental limit theorems work. ‘Stationary’ means that probabilities do not change in time, and ‘ergodic’ that cross-sectional means converge to long-time averages. Between pieces it is necessary to invoke phase change formalism, a ‘biological’ renormalization that generalizes Wilson’s (1971) approach to physical phase transition (Wallace, 2005a).

Shannon uncertainties *H*(…) are cross-sectional law-of-large-numbers sums of the form 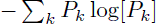, where the *P_k_* constitute a probability distribution. See Cover and Thomas (2006), Ash (1990), or Khinchin (1957) for the standard details.

For cognitive systems, an equivalence class algebra can be constructed by choosing different origin points *a*_0_, and defining the equivalence of two states *a_m_*, *a_n_* by the existence of high probability meaningful paths connecting them to the same origin point. Disjoint partition by equivalence class, analogous to orbit equivalence classes for a dynamical system, defines the vertices of a network of cognitive dual languages that interact to actually constitute the system of interest. Each vertex then represents a different information source dual to a cognitive process. This is not a representation of a network of interacting physical systems as such, in the sense of network systems biology (e.g., Arrell and Terzic, 2010). It is an abstract set of languages dual to the set of cognitive processes of interest, that may become linked into higher order structures.

Topology, in the 20th century, became an object of algebraic study, so-called algebraic topology, via the fundamental underlying symmetries of geometric spaces. Rotations, mirror transformations, simple (‘affine’) displacements, and the like, uniquely characterize topological spaces, and the networks inherent to cognitive phenomena having dual information sources also have complex underlying symmetries: characterization via equivalence classes defines a groupoid, an extension of the idea of a symmetry group, as summarized by Brown (1987) and Weinstein (1996). Linkages across this set of languages occur via the groupoid generalization of Landau’s spontaneous symmetry breaking arguments that will be used below (Landau and Lifshitz, 2007; Pettini, 2007). See the Mathematical Appendix for a brief summary of basic material on groupoids.

## 4 Environment as an information source

Multifactorial cognitive and behavioral systems interact with, affect, and are affected by, embedding environments that ‘remember’ interaction by various mechanisms. It is possible to reexpress environmental dynamics in terms of a grammar and syntax that represent the output of an information source – another generalized language.

Some examples:

1. The turn-of-the seasons in a temperate climate, for many ecosystems, looks remarkably the same year after year: the ice melts, the migrating birds return, the trees bud, the grass grows, plants and animals reproduce, high summer arrives, the foliage turns, the birds leave, frost, snow, the rivers freeze, and so on.
2. Human interactions take place within fairly well defined social, cultural, and historical constraints, depending on context: birthday party behaviors are not the same as cocktail party behaviors in a particular social set, but both will be characteristic.
3. Gene expression during development is highly patterned by embedding environmental context via ‘norms of reaction’ (e.g., Wallace and Wallace, 2010).

Suppose it possible to coarse-grain the generalized ‘ecosystem’ at time *t*, in the sense of symbolic dynamics (e.g., Beck and Schlogl, 1993) according to some appropriate partition of the phase space in which each division *A_j_* represent a particular range of numbers of each possible fundamental actor in the generalized ecosystem, along with associated larger system parameters. What is of particular interest is the set of longitudinal paths, system statements, in a sense, of the form *x*(*n*) = *A*_0_, *A*_1_,…, *A_n_* defined in terms of some natural time unit of the system. Thus *n* corresponds to an again appropriate characteristic time unit *T*, so that *t* = *T*, 2*T*,…, *nT*.

Again, the central interest is in serial correlations along paths.

Let *N*(*n*) be the number of possible paths of length *n* that are consistent with the underlying grammar and syntax of the appropriately coarsegrained embedding ecosystem, in a large sense. As above, the fundamental assumptions are that – for this chosen coarse-graining – *N*(*n*), the number of possible grammatical paths, is much smaller than the total number of paths possible, and that, in the limit of (relatively) large *n*, *H* = lim_*n*→∞_ log[*N*(*n*)]/*n* both exists and is independent of path.

These conditions represent a parallel with parametric statistics systems for which the assumptions are not true will require specialized approaches.

Nonetheless, not all possible ecosystem coarse-grainings are likely to work, and different such divisions, even when appropriate, might well lead to different descriptive quasi-languages for the ecosystem of interest. Thus, empirical identification of relevant coarse-grainings for which this theory will work may represent a difficult scientific problem.

Given an appropriately chosen coarse-graining, define joint and conditional probabilities for different ecosystem paths, having the form *P*(*A*_0_, *A*_1_,…, *A*_*n*_), *P*(*A*_*n*_|*A*_0_,…, *A*_*n*−1_), such that appropriate joint and conditional Shannon uncertainties can be defined on them that satisfy equation (2).

Taking the definitions of Shannon uncertainties as above, and arguing backwards from the latter two parts of equation (2), it is indeed possible to recover the first, and divide the set of all possible ecosystem temporal paths into two subsets, one very small, containing the grammatically correct, and hence highly probable paths, that we will call ‘meaningful’, and a much larger set of vanishingly low probability.

## 5 Body dynamics as an information source

Body movement is inherently constrained by evolutionary Bauplan: snakes do not brachiate, humans cannot (easily) scratch their ears with their hind legs, fish do not breathe air, nor mammals water. This is so evident that one simply does not think about it. Nonetheless, teaching a human to walk and talk, a bird to fly, or a lion to hunt, in spite of evolution, are arduous enterprises that take considerable attention from parents or even larger social groupings. Given the basic bodyplan of head and four limbs, or two feet and wings, or of a limbless spine, the essential point is that not all motions are possible. Bauplan imposes limits on dynamics. That is, if we coarsegrain motions, perhaps using some form of the standard methods for choreography transcription appropriate to the organism under study, we see immediately that not all ‘statements’ possible using the dance symbols have the same probability. That is, there will inevitably be a grammar and syntax to observed body-based behaviors imposed by evolutionary bauplan. Sequences of symbols, say of length n, representing observed motions can be segregated into two sets, the first, and vastly larger, consisting of meaningless sequences (like humans scratching their ears with their feet) that have vanishingly small probability as *n* → ∞ The other set, consistent with underlying bauplan grammar and syntax, can be viewed as the output of an information source, in precisely the manner of the previous two sections, in first approximation following the relations of equation (2).

## 6 Interacting information sources

Given a set of information sources that are linked to solve a problem, in the sense of Wilson and Golonka (2013), the ‘no free lunch’ theorem (English, 1996; Wolpert and Macready, 1995, 1997) extends a network theory-based theory (e.g., Arrell and Terzic, 2010). Wolpert and Macready show there exists no generally superior computational function optimizer. That is, there is no ‘free lunch’ in the sense that an optimizer pays for superior performance on some functions with inferior performance on others gains and losses balance precisely, and all optimizers have identical average performance. In sum, an optimizer has to pay for its superiority on one subset of functions with inferiority on the complementary subset.

This result is well-known using another description. Shannon (1959) recognized a powerful duality between the properties of an information source with a distortion measure and those of a channel. This duality is enhanced if we consider channels in which there is a cost associated with the different letters. Solving this problem corresponds to finding a source that is right for the channel and the desired cost. Evaluating the rate distortion function for a source corresponds to finding a channel that is just right for the source and allowed distortion level.

Yet another approach to the same result is the through the ‘tuning theorem’ (Wallace, 2005a, Sec. 2.2), which inverts the Shannon Coding Theorem by noting that, formally, one can view the channel as ‘transmitted’ by the signal. Then a dual channel capacity can be defined in terms of the channel probability distribution that maximizes information transmission assuming a fixed message probability distribution.

From the no free lunch argument, Shannon’s insight, or the ‘tuning theorem’, it becomes clear that different challenges facing any cognitive system, distributed collection of them, or interacting set of other information sources, that constitute an organism or automaton, must be met by different arrangements of cooperating modules represented as information sources.

It is possible to make a very abstract picture of this phenomenon based on the network of linkages between the information sources dual to the individual ‘unconscious’ cognitive modules (UCM), and those of related information sources with which they interact. That is, a shifting, task-mapped, network of information sources is continually reexpressed: given two distinct problems classes confronting the organism or automaton, there must be two different wirings of the information sources, including those dual to the available UCM, with the network graph edges measured by the amount of information crosstalk between sets of nodes representing the different sources.

Thus fully embodied systems, in the sense of Wilson and Golonka (2013), involve interaction between very general sets of information sources assembled into a ‘task-specific device’ in the sense of Bingham (1988) that is necessarily highly tunable. This mechanism represents a broad evolutionary generalization of the ‘shifting spotlight’ characterizing the global neuronal workspace model of consciousness (Wallace, 2005a).

The mutual information measure of cross-talk is not inherently fixed, but can continuously vary in magnitude. This suggests a parameterized renormalization: the modular network structure linked by crosstalk has a topology depending on the degree of interaction of interest.

Define an interaction parameter *ω*, a real positive number, and look at geometric structures defined in terms of linkages set to zero if mutual information is less than, and ‘renormalized’ to unity if greater than, *ω*. Any given *ω* will define a regime of giant components of network elements linked by mutual information greater than or equal to it.

Now invert the argument: a given topology for the giant component will, in turn, define some critical value, *ω_C_*, so that network elements interacting by mutual information less than that value will be unable to participate, i.e., will be locked out and not be consciously or otherwise perceived. See Wallace (2005a, 2012) for details. Thus *ω* is a tunable, syntactically-dependent, detection limit that depends critically on the instantaneous topology of the giant component of linked information sources defining the analog to a global broadcast of consciousness. That topology is the basic tunable syntactic filter across the underlying modular structure, and variation in *ω* is only one aspect of more general topological properties that can be described in terms of index theorems, where far more general analytic constraints can become closely linked to the topological structure and dynamics of underlying networks, and, in fact, can stand in place of them (Atyah and Singer, 1963; Hazewinkel, 2002).

## 7 Simple regulation

Continuing the formal theory, information sources are often not independent, but are correlated, so that a joint information source – representing, for example, the interaction between brain, body, and the environment – can be defined having the properties

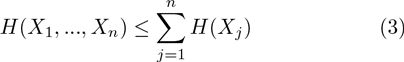

with equality only for isolated, independent information streams.

This is the information chain rule (Cover and Thomas, 2006), and has implications for free energy consumption in regulation and control of embodied cognitive processes. Feynman (2000) describes how information and free energy have an inherent duality, defining information precisely as the free energy needed to erase a message. The argument is quite direct, and it is easy to design an idealized machine that turns the information within a message directly into usable work – free energy. Information is a form of free energy and the construction and transmission of information within living things – the physical instantiation of information – consumes considerable free energy, with inevitable – and massive – losses via the second law of thermodynamics.

Suppose an intensity of available free energy is associated with each defined joint and individual information source *H*(*X,Y*), *H*(*X*), *H*(*Y*), e.g., rates *M_X,Y_*, *M_X_*, *M*_*Y*_.

Although information is a form of free energy, there is necessarily a massive entropic loss in its actual expression, so that the probability distribution of a source uncertainty *H* might be written in Gibbs form as

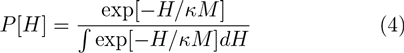

assuming *к* is very small.

To first order, then,

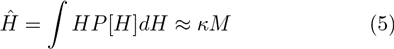

and, using equation (3),

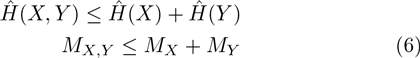

Thus, as a consequence of the information chain rule, allowing crosstalk consumes a lower rate of free energy than isolating information sources. That is, in general, it takes more free energy – higher total cost – to isolate a set of cognitive phenomena and an embedding environment than it does to allow them to engage in crosstalk (Wallace, 2012).

Hence, at the free energy expense of supporting two information sources, – *X* and *Y* together – it is possible to catalyze a set of joint paths defined by their joint information source. In consequence, given a cognitive module (or set of them) having an associated information source *H*(…), an external information source *Y* – the embedding environment – can catalyze the joint paths associated with the joint information source *H*(…, *Y*) so that a particular chosen developmental or behavioral pathway – in a large sense – has the lowest relative free energy.

At the expense of larger global free information expenditure – maintaining two (or more) information sources with their often considerable entropic losses instead of one – the system can feed, in a sense, the generalized physiology of a Maxwell’s Demon, doing work so that environmental signals can direct system cognitive response, thus *locally* reducing uncertainty at the expense of larger global entropy production.

Given a cognitive biological system characterized by an information source *X*, in the context of – for humans – an explicitly, slowly-changing, cultural ‘environmental’ information source *Y*, we will be particularly interested in the joint source uncertainty defined as *H*(*X,Y*), and next examine some details of how such a mutually embedded system might operate in real time, focusing on the role of rapidly-changing feedback information, via the Data Rate Theorem.

## 8 Phase transition

A fundamental homology between the information source uncertainty dual to a cognitive process and the free energy density of a physical system arises, in part, from the formal similarity between their definitions in the asymptotic limit. Information source uncertainty can be defined as in the first part of equation (2). This is quite analogous to the free energy density of a physical system in terms of the thermodynamic limit of infinite volume (e.g., Wilson, 1971; Wallace, 2005a). Feynman (2000) provides a series of physical examples, based on Bennett’s (1988) work, where this homology is an identity, at least for very simple systems. Bennett argues, in terms of idealized irreducibly elementary computing machines, that the information contained in a message can be viewed as the work saved by not needing to recompute what has been transmitted.

It is possible to model a cognitive system interacting with an embedding environment using a simple extension of the language-of-cognition approach above. Recall that cognitive processes can be formally associated with information sources, and how a formal equivalence class algebra can be constructed for a complicated cognitive system by choosing different origin points in a particular abstract ‘space’ and defining the equivalence of two states by the existence of a high probability meaningful path connecting each of them to some defined origin point within that space.

Recall that disjoint partition by equivalence class is analogous to orbit equivalence relations for dynamical systems, and defines the vertices of a network of cognitive dual languages available to the system: each vertex represents a different information source dual to a cognitive process. The structure creates a large groupoid, with each orbit corresponding to a transitive groupoid whose disjoint union is the full groupoid, and each subgroupoid associated with its own dual information source. Larger groupoids will, in general, have ‘richer’ dual information sources than smaller.

We can now begin to examine the relation between system cognition and the feedback of information from the rapidly-changing real-time (as opposed to a slow-time cultural or other) environment, ℋ, in the sense of equation (1).

With each subgroupoid *G_i_* of the (large) cognitive groupoid we can associate a joint information source uncertainty 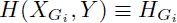, where *X* is the dual information source of the cognitive phenomenon of interest, and *Y* that of the embedding environmental context – largely defined, for humans, in terms of culture and path-dependent historical trajectory.

Recall also that real time dynamic responses of a cognitive system can be represented by high probability paths connecting ‘initial’ multivariate states to ‘final’ configurations, across a great variety of beginning and end points. This creates a similar variety of groupoid classifications and associated dual cognitive processes in which the equivalence of two states is defined by linkages to the same beginning and end states. Thus, we will show, it becomes possible to construct a ‘groupoid free energy’ driven by the quality of rapidly-changing, real-time information coming from the embedding ecosystem, represented by the information rate ℋ, taken as a temperature analog.

For humans in particular, ℋ is an embedding context for the underlying cognitive processes of interest, here the tunable, shifting, global broadcasts of consciousness as embedded in, and regulated by, culture. The argument-by-abduction from physical theory is, then, that ℋ constitutes a kind of thermal bath for the processes of culturally-channeled cognition. Thus we can, in analogy with the standard approach from physics (Pettini, 2007; Landau and Lifshitz, 2007) construct a Morse Function by writing a pseudo-probability for the jointly-defined information sources 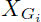, *Y* having source uncertainty 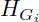 as

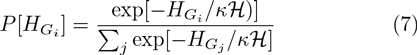

where *к* is an appropriate dimensionless constant characteristic of the particular system. The sum is over all possible sub-groupiods of the largest available symmetry groupoid. Again, compound sources, formed by the (tunable, shifting) union of underlying transitive groupoids, being more complex, will have higher free-energy-density equivalents than those of the base transitive groupoids.

A possible Morse Function for invocation of Pettini’s topological hypothesis or Landau’s spontaneous symmetry breaking is then a ‘groupoid free energy’ *F* defined by

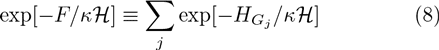

It is possible, using the free energy-analog *F*, to apply Landau’s spontaneous symmetry breaking arguments, and Pettini’s topological hypothesis, to the groupoid associated with the set of dual information sources.

Many other Morse Functions might be constructed here, for example based on representations of the cognitive groupoid(s). The resulting qualitative picture would not be significantly different. We will return to this argument below.

Again, Landau’s and Pettini’s insights regarding phase transitions in physical systems were that certain critical phenomena take place in the context of a significant alteration in symmetry, with one phase being far more symmetric than the other (Landau and Lifshitz, 2007; Pettini, 2007). A symmetry is lost in the transition – spontaneous symmetry breaking. The greatest possible set of symmetries in a physical system is that of the Hamiltonian describing its energy states. Usually states accessible at lower temperatures will lack the symmetries available at higher temperatures, so that the lower temperature phase is less symmetric: The randomization of higher temperatures ensures that higher symmetry/energy states will then be accessible to the system. The shift between symmetries is highly punctuated in the temperature index.

The essential point is that decline in the richness of real-time environmental feedback ℋ, or in the ability of that feed-back to influence response, as indexed by *κ*, can lead to punctuated decline in the complexity of cognitive process within the entity of interest, according to this model.

This permits a Landau-analog phase transition analysis in which the quality of incoming information from the embedding ecosystem – feedback – serves to raise or lower the possible richness of an organism’s cognitive response to patterns of challenge. If *κ*ℋ is relatively large – a rich and varied real-time environment, as perceived by the organism – then there are many possible cognitive responses. If, however, noise or simple constraint limit the magnitude of *κ*ℋ, then behavior collapses in a highly punctuated manner to a kind of ground state in which only limited responses are possible, represented by a simplified cognitive groupoid structure.

Certain details of such information phase transitions can be calculated using ‘biological’ renormalization methods (Wallace, 2005a, Section 4.2) analogous to those used in the determination of physical phase transition universality classes (Wilson, 1971).

These results represent a significant generalization of the Data Rate Theorem, as expressed in equation (1).

Consider, next, an inverse order parameter defined in terms of a conscious attention index, a nonnegative real number *R*. Thus *R* would be a measure of the attention given to the signal defining ℋ. According to the Landau argument, *R* disappears when ℋ ≤ ℋ*_C_*, for some critical value. That is, when ℋ < ℋ*_C_*, there is spontaneous symmetry breaking: only above that value can a global broadcast take place entraining numerous unconscious cognitive submodules, allowing *R* > 0. Below ℋ*_C_*, no global broadcast takes place, and attention is fragmented, or centered elsewhere, so that *R* = 0. A classic Landau order parameter might be constructed as 2/(1 + exp[*aR*]), or 1/[1 + (*aR*)^*n*^], where *a*, *n* ≫ 1.

## 9 Another picture

Here we use the rich vocabulary associated with the stability of stochastic differential equations to model, from another perspective, phase transitions in the composite system of ‘brain/body/environment’ (e.g., Horsthemeke and Lefever, 2006; Van den Broeck et al., 1994, 1997).

Define a ‘symmetry entropy’ based on the Morse Function *F* of equation (8) over a set of structural parameters **Q** = [*Q*_1_,…, *Q_n_*] (that may include ℋ and other information source uncertainties) as the Legendre transform

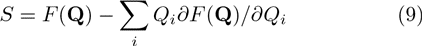

The dynamics of such a system will be driven, at least in first approximation, by Onsager-like nonequilibrium thermodynamics relations having the standard form (de Groot and Mazur, 1984):

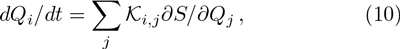

where the *𝒦*_*i,j*_ are appropriate empirical parameters and *t* is the time. A biological system involving the transmission of information may, or may not, have local time reversibility: in English, for example, the string ‘eht’ has a much lower probability than ‘the’. Without microreversibility, *𝒦*_*i,j*_ ≠ *𝒦*_*j,i*_.

Since, however, biological systems are quintessentially noisy, a more fitting approach is through a set of stochastic differential equations having the form

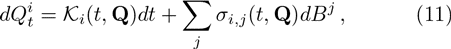

where the *𝒦*_*i*_ and *σ_i,j_* are appropriate functions, and different kinds of ‘noise’ *dB^j^* will have particular kinds of quadratic variation affecting dynamics (Protter, 1990).

Several important dynamics become evident:

1. Setting the expectation of equation (11) equal to zero and solving for stationary points gives attractor states since the noise terms preclude unstable equilibria. Obtaining this result, however, requires some further development.
2. This system may converge to limit cycle or pseudo-random ‘strange attractor’ behaviors similar to thrashing in which the system seems to chase its tail endlessly within a limited venue – a kind of ‘Red Queen’ pathology.
3. What is converged to in both cases is not a simple state or limit cycle of states. Rather it is an equivalence class, or set of them, of highly dynamic modes coupled by mutual interaction through crosstalk and other interactions. Thus ‘stability’ in this structure represents particular patterns of ongoing dynamics rather than some identifiable static configuration or ‘answer’. These are, then, quasi-stationary nonequlibrium states.
4. Applying Ito’s chain rule for stochastic differential equations to the 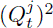 and taking expectations allows calculation of variances. These may depend very powerfully on a system’s defining structural constants, leading to significant instabilities depending on the magnitudes of the *Q*_*i*_, as in the Data Rate Theorem (Khasminskii, 2012).
5. Following the arguments of Champagnat et al. (2006), this is very much a coevolutionary structure, where fundamental dynamics are determined by the feedback between internal and external.

In particular, setting the expectation of equation (11) to zero generates an index theorem (Hazewinkel, 2002) in the sense of Atiah and Singer (1963), that is, an expression that relates analytic results, the solutions of the equations, to underlying topological structure, the eigenmodes of a complicated geometric operator whose groupoid spectrum represents symmetries of the possible changes that must take place for a global workspace to become activated.

Consider, now, the attention measure, *R*, above. Suppose, once triggered, the reverberation of cognitive attention to an incoming signal is explosively self-dynamic – ‘reentrant’ – but that the recognition rate is determined by the magnitude of of the signal κℋ, and affected by noise, so that

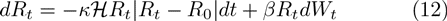

where *dW*_*t*_ represents white noise, and all constants are positive. At steady state, the expectation of equation (12) – the mean attention level – is either zero or the canonical excitation level *R*_0_.

But Wilson (1971) invokes fluctuation at all scales as the essential characteristic of physical phase transition, with invariance under renormalization defining universality classes. Criticality in biological or other cognitive systems is not likely to be as easily classified, e.g., Wallace (2005a, Section 4.2), but certainly failure to have a second moment seems a good analog to Wilson’s instability criterion. As discussed above, analogous results relating phase transitions to noise in stochastic differential equation models are widely described in the physics literature.

To calculate the second moment in *R*, now invoke the Ito chain rule, letting 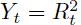. Then

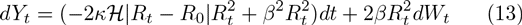

where 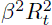 in the *dt* term is the Ito correction due to noise. Again taking the expectation at steady state, no second moment can exist unless the expectation of 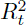 is greater than or equal to zero, giving the condition

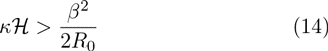

Thus, in consonance with the direct phase transition arguments in ℋ, there is a minimum signal level necessary to support a self-dynamic attention state, in this model. The higher the ‘noise’ – and the weaker the strength of the excited state – the greater the needed environmental signal strength to trigger punctuated ‘reentrant’ attention dynamics.

This result, analogous to equation (1), has evident implications for the quality of attention states in the context of environmental interaction.

## 10 Large deviations

As Champagnat et al. (2006) describe, shifts between the quasi-steady states of a coevolutionary system like that of equation (11) can be addressed by the large deviations formalism. The dynamics of drift away from trajectories predicted by the canonical equations can be investigated by considering the asymptotic of the probability of ‘rare events’ for the sample paths of the diffusion.

‘Rare events’ are the diffusion paths drifting far away from the direct solutions of the canonical equation. The probability of such rare events is governed by a large deviation principle, driven by a ‘rate function’ ℐ that can be expressed in terms of the parameters of the diffusion.

This result can be used to study long-time behavior of the diffusion process when there are multiple attractive singularities. Under proper conditions, the most likely path followed by the diffusion when exiting a basin of attraction is the one minimizing the rate function ℐ over all the appropriate trajectories.

An essential fact of large deviations theory is that the rate function ℐ almost always has the canonical form

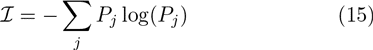

for some probability distribution (Dembo and Zeitouni, 1998).

The argument relates to equation (11), now seen as subject to large deviations that can themselves be described as the output of an information source (or sources), say *L*_*D*_, defining ℐ, driving *Q^j^*-parameters that can trigger punctuated shifts between quasi-steady state topological modes of interacting cognitive submodules.

It should be clear that both internal and feedback signals, and independent, externally-imposed perturbations associated with the source uncertainty ℐ, can cause such transitions in a highly punctuated manner. Some impacts may, in such a coevolutionary system, be highly pathological over a developmental trajectory, necessitating higher order regulatory system counterinterventions over a subsequent trajectory.

Similar ideas are now common in systems biology (e.g., Kitano 2004).

## 11 Canonical failure modes

An information source defining a large deviations rate function ℐ in equation (15) can also represent input from ‘unexpected or unexplained internal dynamics’ (UUID) unrelated to external perturbation. Such UUID will always be possible in sufficiently large cognitive systems, since crosstalk between cognitive submodules is inevitable, and any possible critical value will be exceeded if the structure is large enough or is driven hard enough. This suggests that, as Nunney (1999) describes for cancer, large-scale cognitive systems must be embedded in powerful regulatory structures over the life course. Wallace (2005b), in fact, examines a ‘cancer model’ of regulatory failure for mental disorders.

More specifically, the arguments leading to equations (7) and (8) could be reexpressed using a joint information source

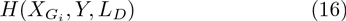

providing a more complete picture of large-scale cognitive dynamics in the presence of embedding regulatory systems, or of sporadic external ‘therapeutic’ interventions. However, the joint information source of equation (16) now represents a de-facto distributed cognition involving interpenetration between both the underlying embodied cognitive process and its similarly embodied regulatory machinery.

That is, we can now define a composite Morse Function of embodied cognition-and-regulation, *F*_*ECR*_, as

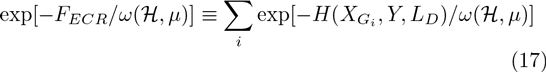

where *ω* (ℋ, *μ*) is a monotonic increasing function of both the data rate ℋ and of the ‘richness’ of the internal cognitive function defined by an internal – strictly cognitive – network coupling parameter *μ*, a more limited version of the argument in Section 6. Typical examples might include 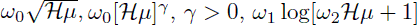, and so on.

More generally, 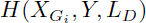 in equation (17) could probably be replaced by the norm

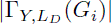

for appropriately chosen representations Γ of the underlying cognitive-defined groupoid, in the sense of Bos (2007) and Buneci (2003). That is, many Morse Functions similarly parameterized by the monotonic functions *ω* (*ℋ*, *μ*) are possible, with the underlying topology, in the sense of Pettini, itself more subtly parameterized, in a way, by the information sources *Y* and *L*_*D*_.

Applying Pettini’s topological hypothesis to the chosen Morse Function, reduction of either ℋ or *μ*, or both, can trigger a ‘ground state collapse’ representing a phase transition to a less (groupoid) symmetric ‘frozen’ state. In higher organisms, which must generally function under real-time constraints, elaborate secondary back-up systems have evolved to take over behavioral control under such conditions. These typically range across basic emotional, as well as hypothalamic-pituitary-adrenal (HPA) and hypothalamic-pituitary-thyroid (HPT) axis, responses (e.g., Wallace, 2005a, 2012, 2013; Wallace and Fullilove, 2008). Dysfunctions of these systems are implicated across a vast spectrum of common, and usually comorbid, mental and physical disorders (e.g., Wallace, 2005a, b; Wallace and Wallace, 2010, 2013).

Given the inability of some half-billion years of evolutionary selection pressures to successfully overcome such challenges – comorbid mental and physical disorders before senescence remain rampant in human populations – it seems unlikely that automatons or man-machine cockpits designed for the control of critical real-time systems can avoid ground-state collapse and other critical failure modes, if naively deployed (e.g., Hawley, 2006, 2008). Indeed, the conundrum of ‘robot emotions’ has already engendered considerable study (e.g., Fellous and Arbib, 2005).

## 12 Discussion and conclusions

Bernard Baars’ global workspace model of consciousness (Baars, 1988; Baars et al., 2013; Wallace, 2005a) posits a ‘theater spotlight’ involving the recruitment of unconscious cognitive modules of the brain into a temporary, tunable, general broadcast fueled by crosstalk that allows formation of the shifting coalitions needed to address real-time problems facing a higher organism. Similar exaptations of crosstalk between cognitive modules at smaller scales have been recognized in wound healing, the immune system, and so on (Wallace, 2012; 2014). Newly-developed views of embodied cognition envision that phenomenon as analogous, that is, as the temporary assembly of interacting modules from brain, body, and environment to address real-time problems facing an organism (or a machine). This is likewise a dynamic process that sees many available information sources – not limited to those dual to cognitive brain or internal physiological modules – again linked by crosstalk into a tunable real-time phenomenon that might well be characterized as a generalized consciousness.

Here, we have made formal use of the Data Rate Theorem in exploring the dynamics of such an embodied cognition, and of a necessarily related embodied regulation. These, according to theory, inevitably involve a synergistic interpenetration among nested sets of actors, represented here as information sources. These may include dual sources to internal cognitive modules, body bauplan, environmental information, language, culture, and so on.

To summarize, two factors determine the possible range of real-time cognitive response, in the simplest version of the model. These are the magnitude of of the environmental feedback signal *к*ℋ and the inherent structural richness of the cognitive groupoid defining *F*. If that richness is lacking – if the possibility of internal *μ*-connections is limited – then even very high levels of ℋ may not be adequate to activate appropriate behavioral responses to important real-time feedback signals, following the argument of equation (17).

Cognition and regulation must, then, be viewed as interacting gestalt processes, involving not just an atomized individual (or, taking an even more limited ‘NIMH’ perspective, just the brain of that individual), but the individual in a rich context that must include both the body that acts on the environment, and the environment that reacts on body and brain.

The large deviations analysis suggests that cognitive function also occurs in the context, not only of a powerful environmental embedding, but of a specific regulatory milieu: there can be no cognition without regulation. The ‘stream of generalized consciousness’ represented by embodied cognition must be contained within regulatory riverbanks.

For humans, and many other animal species (e.g., Avital and Jabolonka, 2000), this picture must be expanded by another layer of information sources: as the evolutionary anthropologist Robert Boyd has expressed it, ‘Culture is as much a part of human biology as the enamel on our teeth’. Thus, for humans, the schematic hierarchy of interacting information sources becomes

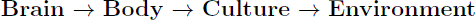

Current theoretical schema for embodied cognition omit the critical level of cultural modulation.

Mental disorders, from this perspective, can be represented as a synergistic dysfunction of internal function and regulatory milieu, most simply characterized by the interaction between the driving parameters *μ* and ℋ.

We have, in a way, extended the criticisms of Bennett and Hacker (2003) who explored the mereological fallacy of a decontextualization that attributes to ‘the brain’ what is the province of the whole individual. Here, we argue that the ‘whole individual’ involves essential interactions with embedding environmental and regulatory settings that, for humans, must include cultural heritage. Real-time automatons and man-machine cockpits, currently expected to function without such an embedding and pervasive regulatory milieu, may face serious stability challenges, particularly subject to a ground state collapse in which ‘all possible targets are enemies’.

## 13 Mathematical appendix

### 13.1 Morse Theory

Morse Theory explores relations between analytic behavior of a function – the location and character of its critical points – and the underlying topology of the manifold on which the function is defined. We are interested in a number of such functions, for example information source uncertainty on a parameter space and possible iterations involving parameter manifolds determining critical behavior. An example might be the sudden onset of a giant component. These can be reformulated from a Morse Theory perspective (Pettini, 2007).

The basic idea of Morse Theory is to examine an *n*-dimensional manifold *M* as decomposed into level sets of some function 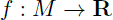 where **R** is the set of real numbers. The *a*-level set of *f* is defined as

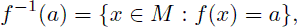

the set of all points in *M* with *f*(*x*) = *a*. If *M* is compact, then the whole manifold can be decomposed into such slices in a canonical fashion between two limits, defined by the minimum and maximum of *f* on *M*. Let the part of *M* below *a* be defined as

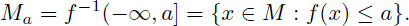

These sets describe the whole manifold as *a* varies between the minimum and maximum of *f*.

Morse functions are defined as a particular set of smooth functions 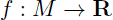 as follows. Suppose a function *f* has a critical point *x_c_*, so that the derivative *df*(*x_c_*) = 0, with critical value *f*(*x_c_*). Then, *f* is a Morse function if its critical points are nondegenerate in the sense that the Hessian matrix of second derivatives at *x_c_*, whose elements, in terms of local coordinates are

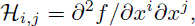

has rank *n*, which means that it has only nonzero eigenvalues, so that there are no lines or surfaces of critical points and, ultimately, critical points are isolated.

The index of the critical point is the number of negative eigenvalues of ℋ at *x_c_*.

A level set *f*^−1^(a) of *f* is called a critical level if *a* is a critical value of *f*, that is, if there is at least one critical point *x_c_* ∈ *f*^−1^(*a*).

Again following Pettini (2007), the essential results of Morse Theory are:

1. If an interval [*a*, *b*] contains no critical values of *f*, then the topology of *f*^−1^ [*a*, *v*] does not change for any *v* ∈ (*a*, *b*]. Importantly, the result is valid even if *f* is not a Morse function, but only a smooth function.
2. If the interval [*a, b*] contains critical values, the topology of *f*^−1^ [*a, v*] changes in a manner determined by the properties of the matrix *H* at the critical points.
3. If *f* : *M* → **R** is a Morse function, the set of all the critical points of *f* is a discrete subset of *M*, i.e., critical points are isolated. This is Sard’s Theorem.
4. If *f* : *M* → **R** is a Morse function, with *M* compact, then on a finite interval [*a, b*] ⊂ **R**, there is only a finite number of critical points *p* of f such that *f* (*p*) ∈ [*a, b*]. The set of critical values of *f* is a discrete set of **R**.
5. For any differentiable manifold *M*, the set of Morse functions on *M* is an open dense set in the set of real functions of *M* of differentiability class *r* for 0 ≤ *r* ≤ ∞.
6. Some topological invariants of *M*, that is, quantities that are the same for all the manifolds that have the same topology as *M*, can be estimated and sometimes computed exactly once all the critical points of f are known: let the Morse numbers *μ*_*i*_(*i* = 0,…, *m*) of a function *f* on *M* be the number of critical points of *f* of index *i*, (the number of negative eigenvalues of *H*). The Euler characteristic of the complicated manifold *M* can be expressed as the alternating sum of the Morse numbers of any Morse function on *M*,

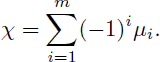

The Euler characteristic reduces, in the case of a simple polyhedron, to

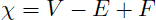

where *V*, *E*, and *F* are the numbers of vertices, edges, and faces in the polyhedron.
7. Another important theorem states that, if the interval [*a, b*] contains a critical value of *f* with a single critical point *x_c_*, then the topology of the set *M*_*b*_ defined above differs from that of *M*_*a*_ in a way which is determined by the index, *i*, of the critical point. Then *M*_*b*_ is homeomorphic to the manifold obtained from attaching to *M*_*a*_ an *i*-handle, i.e., the direct product of an *i*-disk and an (*m* – *i*)-disk.

Pettini (2007) and Matsumoto (2002) contain details and further references.

### 13.2 Groupoids

A groupoid, *G*, is defined by a base set *A* upon which some mapping – a morphism – can be defined. Note that not all possible pairs of states (*a*_*j*_, *a*_*k*_) in the base set *A* can be connected by such a morphism. Those that can define the groupoid element, a morphism *g* = (*a*_*j*_, *a*_*k*_) having the natural inverse *g*^−1^ = (*a*_*k*_, *a*_*j*_). Given such a pairing, it is possible to define ‘natural’ end-point maps *α*(*g*) = *a*_*j*_, *β*(*g*) = *a*_*k*_ from the set of morphisms *G* into *A*, and a formally associative product in the groupoid *g*_1_*g*_2_ provided *α*(*g*_1_*g*_2_) = *α*(*g*_1_), *β*(*g*_1_*g*_2_) = *β*e(*g*_2_), and *β*(*g*_1_) = *α*(*g*_2_). Then, the product is defined, and associative, (*g*_1_*g*_2_)*g*_3_ = *g*_1_(*g*_2_*g*_3_). In addition, there are natural left and right identity elements *λ*_*g*_, *ρ*_*g*_ such that *λ*_*g*_*g* = *g* = *gp*_*g*_.

An orbit of the groupoid *G* over *A* is an equivalence class for the relation *a*_*j*_ ∼ *Ga*_*k*_ if and only if there is a groupoid element *g* with *α*(*g*) = *a*_*j*_ and *β*(*g*) = *a*_*k*_. A groupoid is called transitive if it has just one orbit. The transitive groupoids are the building blocks of groupoids in that there is a natural decomposition of the base space of a general groupoid into orbits. Over each orbit there is a transitive groupoid, and the disjoint union of these transitive groupoids is the original groupoid. Conversely, the disjoint union of groupoids is itself a groupoid.

The isotropy group of *a* ∈ *X* consists of those *g* in *G* with *α*(*g*) = *a* = *β*(*g*). These groups prove fundamental to classifying groupoids.

If *G* is any groupoid over *A*, the map (*α*, *β*) : *G* → *A* × *A* is a morphism from *G* to the pair groupoid of *A*. The image of (*α*, *β*) is the orbit equivalence relation 〜 *G*, and the functional kernel is the union of the isotropy groups. If *f* : *X* → *Y* is a function, then the kernel of *f*, *ker*(*f*) = [(*x*_1_, *x*_2_) ∈ *X* × *X* : *f* (*x*_1_) = *f*(*x*_2_)] defines an equivalence relation.

Groupoids may have additional structure. For example, a groupoid *G* is a topological groupoid over a base space *X* if *G* and *X* are topological spaces and *α*, *β* and multiplication are continuous maps.

In essence, a groupoid is a category in which all morphisms have an inverse, here defined in terms of connection to a base point by a meaningful path of an information source dual to a cognitive process.

The morphism (*α*, *β*) suggests another way of looking at groupoids. A groupoid over *A* identifies not only which elements of A are equivalent to one another (isomorphic), but *it also parameterizes the different ways (isomorphisms) in which two elements can be equivalent*, i.e., in our context, all possible information sources dual to some cognitive process. Given the information theoretic characterization of cognition presented above, this produces a full modular cognitive network in a highly natural manner.

